# Experimental mismatch in benchmarking PELSA and LiP-MS

**DOI:** 10.64898/2026.03.24.713688

**Authors:** Chloé Van Leene, Emin Araftpoor, Kris Gevaert

**Affiliations:** VIB UGent Center for Medical Biotechnology, VIB, Ghent, Belgium; Department of Biomolecular Medicine, Ghent University, Ghent, Belgium

## Abstract

Limited proteolysis coupled to mass spectrometry (**LiP-MS**) is a peptide-centric conformational proteomics approach during which a brief incubation with a non-specific protease (e.g., proteinase K) under native conditions generates structural fingerprints that report on treatment-induced conformational changes, which is followed by a tryptic digest under denaturing conditions allowing to read out these fingerprints ^1^. In contrast, the recently introduced peptide-centric local stability assay (**PELSA**) uses a high trypsin-to-substrate ratio under native conditions to release fully tryptic peptides that reflect structural stability upon ligand binding ^2^. In their paper, Li *et al*. compared PELSA and LiP-MS across several benchmarks and reported that PELSA exhibited quantitative sensitivity comparable to or exceeding LiP-MS. Notably, PELSA quantified a 21-fold greater rapamycin-induced change for FKBP1A compared to LiP-MS. Because such claims influence method selection for conformational proteomics, we reanalyzed the publicly deposited datasets underlying these comparisons and assessed the experimental and analytical choices that contributed to the reported effect sizes. Our evaluation indicates that the reported 21-fold difference arises from non-matched experimental conditions and undisclosed data imputation, and that conclusions regarding quantitative superiority or biological interpretability should therefore be treated with caution.

## Differences in experimental design preclude quantitative comparison

Although Li *et al*. present head-to-head comparisons, the PELSA and LiP-MS datasets for rapamycin were not generated under comparable conditions. The PELSA rapamycin experiment was performed following a 30-minute incubation with 2 µM rapamycin at 25 °C and acquired on an Orbitrap Exploris 480 ^2^, whereas the published LiP-MS dataset used a 10-minute incubation with 2 µM rapamycin at room temperature and a Q-Exactive HF instrument. Beyond these experimental differences, the LC-MS acquisition parameters also diverged substantially, including differences in solvent gradient, instrument type, chromatographic setup and DIA scheme ^3^. Furthermore, the DIA searches were processed in different Spectronaut versions (10 vs. 15), which differ substantially in their underlying scoring and processing algorithms (**Table 1**).

**Table 1.** Methodological differences between the published LiP-MS and PELSA datasets that preclude direct quantitative comparison.

|  | PELSA data | LiP-MS dataset <sup>3</sup> |
| --- | --- | --- |
| *Treatment incubation time and temperature* | 25 °C, 30 minutes | Room temperature, 10 minutes |
| *LC metrics* | Micro-flow LC, linear gradient | Nano-flow, segmented gradient |
| *MS instrument* | Orbitrap Exploris 480 | Q-Exactive HF |
| *Sample load* | 10 µg peptide material | 2 µg peptide material |
| *Spectronaut version* | Reported as Spectronaut 15.3 | Processed originally in Spectronaut 10 |

These discrepancies have direct consequences for data interpretation. Ligand-target kinetic studies show that incubation time strongly influences the extent of target engagement, with longer incubations increasing the likelihood of secondary or off-target interactions as equilibrium shifts over time. Accordingly, the difference in ligand incubation time between the two assays (10 min vs. 30 min) may have already altered the number and magnitude of peptide-level responses ^4^. In combination with fundamentally different LC-MS conditions and different sample loads (10 µg vs. 2 µg), all of which affect precursor detectability and quantitative depth, the resulting measurements cannot be correctly compared in a quantitative manner. Because peptide-level fold changes in conformational proteomics depend strongly on acquisition depth, MS platform sensitivity and missingness patterns, comparing effect sizes across such heterogeneous acquisition and processing conditions risks (over-)interpreting technical differences as biological differences.

## Differences in data processing and imputation further confound the comparison

Using the deposited data (PRIDE: PXD034606), we found that the PELSA rapamycin dataset was processed using Spectronaut 18, not Spectronaut 15.3 as stated in the paper. Crucially, all datasets we inspected had undergone imputation, although this was not stated in the manuscript. Biognosys confirmed that Spectronaut versions up to and including 15 impute by default, and the presence of Spectronaut 18 output files indicates that the authors’ final results were generated using defaults that differ from those described.

Imputation is widely used in quantitative proteomics and its influence on effect direction and statistical significance is well known ^5,6^. Claims of method sensitivity should therefore also report the impact of imputation explicitly. In this case, the central quantitative claim, the 21-fold difference, depends on imputed values for precursors that are fully missing in the rapamycin condition. As such, it is not possible to derive a quantitative fold change. Moreover, the reported protein-level effect was inferred from a single strongly changing peptide rather than from the collective behaviour of all FKBP1A peptides, a practice that does not reflect peptide-level consistency and therefore cannot be used to assess method sensitivity.

To quantify the impact of imputation, we reanalyzed the PELSA rapamycin dataset with and without imputation (**Figure 1**). For downstream analysis, we used our DIA-LiPA pipeline ^7^. Without imputation, only one FKBP1A precursor is found significantly altered, and multiple FKBP1A precursors are entirely missing in the rapamycin condition. With imputation, these missing precursors become significant, altering both the statistical output and the sequence-level interpretation.

**Figure 1.**
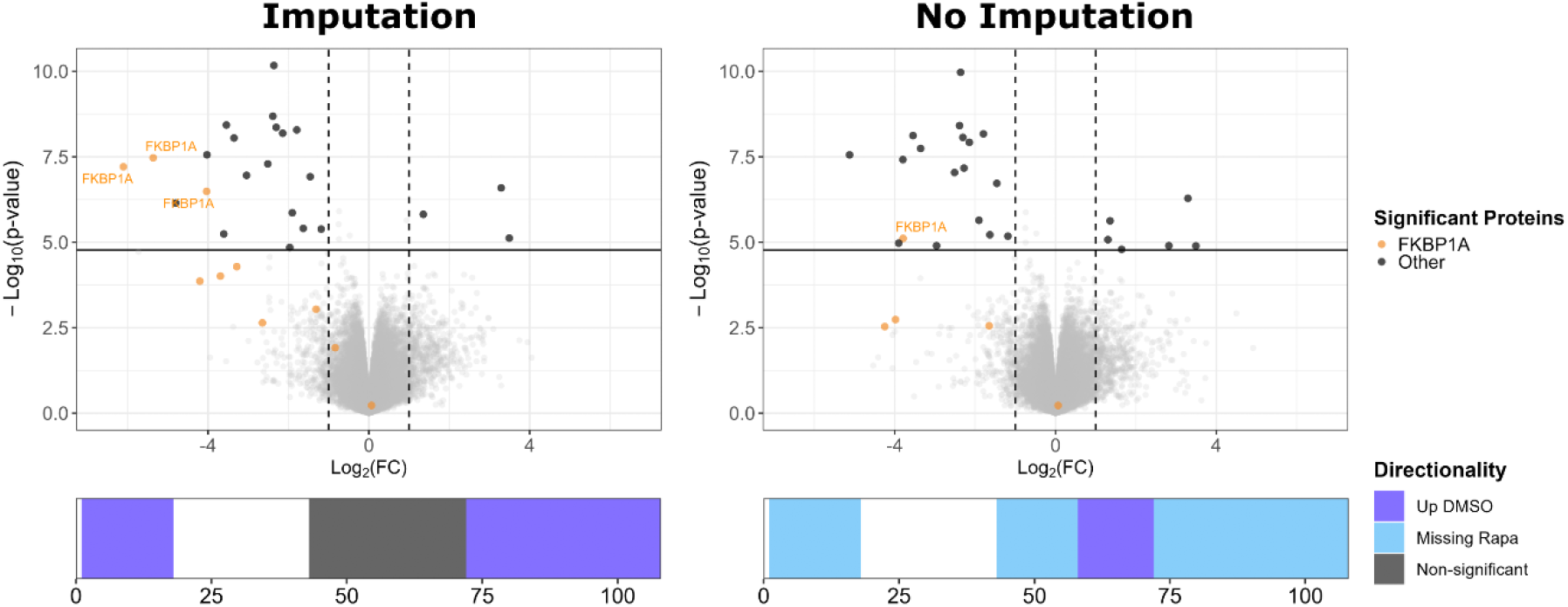
Downstream structural output from PELSA analysis, with (left) and without (right) imputation, analyzed using DIA-LiPA. Volcano plots show precursor-level differential abundance between rapamycin-treated and solvent control samples. Significance thresholds were |log_2_(FC)| ≥ 1 and adjusted p-value ≤ 0.05. FKBP1A precursors are highlighted in orange. Other significant precursors are shown in black, and non-significant precursors in grey. Below each volcano plot, structural barcodes summarize FKBP1A results. Bars represent precursor positions along the protein sequence and are colored by directionality: dark blue (significant and upregulated in solvent-control samples), light blue (missing in rapamycin-treated samples), grey (not significant), and white (not detected). FC, fold change.

A striking example is the 59-72 region of FKBP1A. Without imputation, this region is significantly upregulated in solvent-control samples, consistent with its known location in the rapamycin binding pocket ^8^. After imputation, it no longer appears significant, despite the underlying biology being unchanged. Similar behavior is observed for other FKBP1A regions, demonstrating that imputation can shift structural conclusions.

These observations highlight that missingness patterns, rather than imputed abundances, should be considered when comparing method sensitivity and interpreting protein conformations. Because precursor missingness in conformational proteomics data reflects instrument sensitivity and digestion properties, it may not be assumed to represent structural differences unless validated experimentally by orthogonal experiments.

## Conclusions

PELSA is a valuable addition to the conformational proteomics toolkit: its single-step digestion simplifies sample preparation, produces fully tryptic peptides, and can sensitively detect ligand binding. However, the quantitative comparison between PELSA and LiP-MS presented by Li *et al*. does not permit a reliable assessment of method sensitivity. The datasets differ in incubation times, instrumentation and software versions, and a quantitative comparison between two independently processed datasets that rely on imputed peptide intensities is not methodologically valid. Together, these discrepancies introduce interpretation challenges that preclude a meaningful comparison of sensitivity across methods.

We recommend that future head-to-head evaluations between conformational proteomics methods use matched experimental conditions, report software versions and imputation settings, and consider the presence of missingness in the evaluation of method sensitivity. Transparent reporting of these elements is essential for deriving quantitative conclusions about method sensitivity and ensuring that the field maintains rigorous standards for comparative benchmarking.

## METHODS

The Spectronaut SNE file for the PELSA rapamycin dataset was downloaded from PRIDE (PXD034606). For reanalysis, the dataset was imported into Spectronaut version 18 and imputation was disabled before downstream processing with the DIA-LiPA pipeline. To assess the impact of missing value handling, analyses were performed using the imputed intensities present in the deposited Spectronaut output and using non-imputed values. Precursor-level differential abundance was calculated as log_2_(treated/control), and precursor positions were mapped to the FKBP1A sequence to generate structural barcodes. No protein-level aggregation was performed; all comparisons were kept at the precursor level to avoid representative-peptide selection bias.

## DATA AVAILABILITY

The dataset used was previously published and is publicly available on PRIDE (accession: PXD034606). For this reanalysis, all processed input files used as starting material, including the Spectronaut output files with and without imputation, the FASTA file and the sample annotation, are provided in the Supplementary Information.

## CODE AVAILABILTY

All analyses were conducted using the DIA-LiPA workflow implemented in R. The full analysis structure, including the DIA-LiPA script, all required input files and the generated outputs (volcano plots and structural barcodes), is provided in the Supplementary Information.

## AUTHOR CONTRIBUTIONS

C.V.L. performed the reanalysis and drafted the manuscript. E.A. and K.G. provided feedback on the manuscript and its content.

## NOTES

The authors declare no competing interests.

