## Supplementary material for "Experimental mismatch in benchmarking PELSA and LiP-MS": DIA-LiPA-unpaired_imput.Rmd

````
---
title: "DIA-LiPA (unpaired) reanalysis of original output - imputed"
author: "Chloé Van Leene"
date: "2026-02-23"
#editor: source
output:
  html_document:
  theme: flatly
  code_download: true
  toc: true
  toc_float: true
    #toc-depth: 3
  number_sections: true
    #number-depth: 3

    
---

This script is used to analysis LiP assay data for which a trypsin control (TC) has been carried out. The LiP and TC samples are unpaired/independent. 

# Load packages

We load the `msqrob2` package, along with additional packages for
data manipulation and visualisation.


```{r load_libraries}
suppressPackageStartupMessages({
  library("QFeatures")  
  library("dplyr") 
  library("tidyr")
  library("ggplot2")
  library("ggsci")
  library("ggrepel")
  library("msqrob2")    
  library("stringr")
  library("ExploreModelMatrix")
  library("MsCoreUtils") 
  library("matrixStats")
  library("patchwork")
  library("seqinr")
  library("ComplexHeatmap")
  library("pcaMethods")
  library("scater")
  library("gginnards")
})
```

# Data

## Import Precursor table

We load the output from Spectronaut output file.
1. Define the path to the report file
2. Import the file 

```{r import_data}
precursorFile <- "../input/20260223_rapamycin_rep4_final_Report.tsv"  #1.
library(data.table)
precursors <- fread(precursorFile, sep = "\t") #2.

precursors <- precursors %>%
  mutate(PEP.GroupingKey = paste0(FG.LabeledSequence, ".", FG.Charge))

precursors <- precursors %>%
  rename(
    Run = R.FileName,
    Genes = PG.Genes,
    Protein.Group = PG.ProteinGroups,
    Protein.Names = PG.ProteinNames,
    PG.Q.Value = PG.Qvalue,
    Precursor.Id = PEP.GroupingKey,
    Stripped.Sequence = PEP.StrippedSequence,
    Precursor.Charge = FG.Charge,
    Q.Value = EG.Qvalue,
    Precursor.Quantity = FG.MS2RawQuantity
  )
```


## Import Annotations

We import the sample annotation file
1. Define the path to the sample annotation file
2. Import the file 

```{r}
annotationFile <- "../input/SampleAnnotation.txt" #1.
annotations <- arrow::read_tsv_arrow(annotationFile) |> 
  plyr::rename(c("Run"="runCol")) #2.
```


## Import protein FASTA file 

Import the fasta file with the proteins of the organism that is studied. 
1. Define the path to the report file
2. Read the fasta file

```{r}
fastaFile <- "../input/SP_9606.fasta" #1. 
FASTA <- read.fasta(fastaFile, seqtype = "AA", as.string = T) #2.
```

## Protein mapping

1. Extract the proteins from the protein file. 
2. Calculate the length of the protein

```{r}
# read in human fasta file for protein sequence & length -> to calculate the coverage
mapping <- data.frame(Accession = FASTA |> names() |> str_split_i("\\|", 2),
                      Protein.Sequence = unlist(FASTA)) #1. 
mapping$Protein.Length <- nchar(mapping$Protein.Sequence) #2.
```

## Setup QFeatures object 

We use `Precursor.Quantity` for abundance quantification and `Precursor.Quantity` values below 4 are assumed to be background signal and these entries are filtered. 

```{r}
(qf <- readQFeatures(assayData = precursors |> 
                       filter(Precursor.Quantity > 4), 
                     colData = annotations,
                     quantCols = "Precursor.Quantity",
                     runCol = "Run",
                     fnames = "Precursor.Id"))
```

We redefine Proteotypic proteins. If the protein group consists of one protein it is considered proteotypic, if it consists of multiple proteins the protein group name consists of all proteins separated with a semi colon. 

```{r}
for (i in 1:length(qf))
  rowData(qf[[i]])$Proteotypic <- rowData(qf[[i]])$Protein.Group |> 
             grepl(pattern = ";") |> 
             ifelse(yes = 0, no = 1)
```

## Precursor Filtering 

We apply custom filtering: 

1. q-value threshold of 0.01 for the identification of precursors (`Q.Value`), protein groups (`PG.Q.Value`) and match between run (`Lib.Q.Value`)

2. Remove precursors that could not be mapped, i.e. when `Precursor.Id` column is an empty string.  


```{r filtering}
qf <-  qf |>
  filterFeatures(~ Q.Value <= 0.01 & #1.
                   PG.Q.Value <= 0.01 &  #1. 
                   #Lib.Q.Value <= 0.01 & #1.
                   Precursor.Id != "") #2.
```

## Assay joining

We join the data for the LiP samples.

```{r}
lip_samples <- which(colData(qf)$Pipeline == "LiP")

qf <- joinAssays(
  x = qf,
  i = lip_samples,
  fcol = "Precursor.Id",
  name = "precursors_lip"
  )
```


## Remove highly missing precursors

We first make a function that can extract the smallest group size for an assay, which can be used for filtering. 

1. We get the assay with the corresponding colData
2. We extract the assay data.
3. We check if the data are observed (negate the is.na function).
4. We calculate the group sizes for each row/feature (using matrix multiplication with a dummy matrix for each group). 
5. We caluclate the smallest group size.

```{r}
calculatedSmallestGroupSize <- function(qf, i, facName){
  se <- getWithColData(qf, i = i) #1.
  fac <- colData(se)[[facName]]
  rowData(qf[[i]])$smallestGroupSize <- 
    (se |> 
       assay() |> #2
       Negate(is.na)()  #3
    ) %*% model.matrix(~-1 + fac) |> #4 
    rowMins() #5
  return(qf)
} 
```

We calculate the smallest group size for both assays. 

For the LiP assay we are less stringent as precursors that occur in one condition are informative.  
Here, we keep a precursor as soon as it appears two or more times in the dataset. 

```{r}
qf <- qf |> 
  calculatedSmallestGroupSize(i = "precursors_lip", fac = "Drug") 

nObs <- 2
n <- ncol(qf[["precursors_lip"]])
(qf <- filterNA(qf, i="precursors_lip", pNA=(n-nObs)/n))
```

## Log-transformation 

We log-transform the features in lip assay. 

```{r}
qf <- logTransform(qf, i = "precursors_lip", name="precursors_lip_log")
```

## Normalisation

Then, we apply median normalization based on precursors which are identified in every sample.
We define a helper function for this that 

1. Extracts the assay data 
2. Removes features with missing values
3. Subsequently takes the column wise median to obtain the sample based normalisation factor on the log2 scale.
4. Zero center the normalisation factors
5. Subtract these log2-norm factors from the intensities of each corresponding column of the assay data and store the result in the new assay peptides_norm. (We adopt the sweep function to the peptides_log assay of the spikein QFeatures object with as statistic the log2 normfactor STATS=nf the default function FUN = "-", MARGIN = 2 to substract the column wise log2 norm factor from each entry of the corresponding assay data)

```{r}
medianNormCommonFeatures <- function(qf, i, name="normAssay"){
  norm_factors_lip <- qf[[i]] |> 
    assay() |> #1.
    na.exclude() |> #2.
    colMedians() #3.
  norm_factors_lip <- norm_factors_lip - median(norm_factors_lip)
    qf <- sweep(qf, #4. Subtract log2 norm factor column-by-column (MARGIN = 2)
            MARGIN = 2, 
            STATS = norm_factors_lip, 
            i = i , 
            name = name) 
    return(qf)
}
```

```{r}
qf <- medianNormCommonFeatures(qf, i = "precursors_lip_log", name = "precursors_lip_norm")
```

# QC

We first define helper functions to extract peptide characteristics based on precursor sequence in qfeatures object and protein sequence information in mapping file.  

## Defining helper functions to extract peptide characteristics

We first define a function that classifies peptides into tryptic, semi-tryptic and non-tryptic peptides according to the peptide and protein sequence. 

```{r}
classify_trypticity <- function(peptide, protein, start_pos=NULL) {
  if(is.null(start_pos)) 
    start_pos = ifelse(str_count(protein, peptide)==1, str_locate(protein,peptide) [,"start"], NA)
  pep_len <- sapply(peptide, nchar)
  end_pos <- start_pos + pep_len - 1
 
    # Get surrounding amino acids
  AA_before <- ifelse(start_pos > 1, 
                    substr(protein, start_pos - 1, start_pos - 1), 
                    "")
  AA_after <- ifelse(end_pos < nchar(protein), 
                    substr(protein, end_pos + 1, end_pos + 1), 
                    "")
  AA_first <- substr(peptide, 1, 1)
  AA_last  <- substr(peptide, pep_len, pep_len)

 
  # Classification
  ambiguous <- is.na(protein) | is.na(start_pos)

  nterm_tryptic <- (AA_before %in% c("K", "R", "") | (AA_before == "M" & start_pos==2))
  cterm_tryptic <- (AA_last %in% c("K", "R") | AA_after=="")
  return(
    ifelse(ambiguous, 
         "Ambiguous",
         ifelse(nterm_tryptic & cterm_tryptic,
                "Tryptic",
                ifelse(nterm_tryptic | cterm_tryptic,
                       "Semi-Tryptic",
                       "Non-Tryptic")
                )
    )
  )
 
}
```


Next, we define a function to calculate the protein coverage. 

```{r}
calculate_coverage <- function(start, end, protein_length) {
  require(IRanges)
  #Checks 
  if (is.na(protein_length)) return(NA)
  startEnd <- cbind(start,end) |> na.omit()
  if (nrow(startEnd) <1) return(NA)
  covered <- IRanges(startEnd[,1], startEnd[,2]) |> 
    coverage() |>
    as.logical() |>
    sum()
  return(covered/protein_length)
}
```


## Extract data for QC 

1. We first extract all data that we will use for the QC plots and store it as a long data frame. 
2. We convert it into a data.frame
3. We add the sequence and length of the corresponding protein in the mapping file 
4. We determine the peptide trypticity. 

```{r}
qcObj <- qf[,,c("precursors_lip_norm")] |>
  longForm(colvars = c("Condition",
                       "CondRep",
                       "Drug",
                       "Replicate",
                       "Pipeline"), 
           rowvars= c("Precursor.Id", 
                      "Protein.Group", 
                      "Stripped.Sequence",
                      "Precursor.Charge",
                      "Genes")) |> #1.
  data.frame() |> #2.
  left_join(mapping, by = join_by(Protein.Group == Accession)) |> #3.
  mutate(pep_type = classify_trypticity(Stripped.Sequence,Protein.Sequence)) #4.
```

## ID

We perform QC on the identifications.

```{r}
qcObj |>
  select("Condition", "CondRep", "Pipeline","Precursor.Id", "Protein.Group", "value") |>
  filter(!is.na(value)) |>
  group_by(Condition, CondRep) |>
  summarise(Precursors = length(unique(Precursor.Id)),
            `Protein Groups` = length(unique(Protein.Group))) |>
  pivot_longer(-(1:2),
               names_to = "Feature",
               values_to = "IDs") |>
  ggplot(aes(x = CondRep, y = IDs, fill = Condition)) +
  geom_col() +
  scale_fill_observable() +
  facet_wrap(~Feature,
             scales = "free_y") +
  labs(title = "Precursor and protein group identificiations per sample",
       x = "Sample",
       y = "Identifications") +
  theme_bw() +
  theme(axis.text.x = element_text(angle = 90))
```

## Intensity distributions

We perform QC on the normalised marginal intensity distributions in the different samples. Note that the factor CondRep refers to each sample.

```{r}
qcObj |>
  select("Condition", "CondRep", "Pipeline","value") |>
  filter(!is.na(value)) |>
  ggplot(aes(x = CondRep,
             y = value)) +
    geom_boxplot(aes(fill = Condition)) +
  scale_fill_observable() +
  theme_bw() +
  labs(title = "Normalised precursor quantity distributions per sample",
       x = "Sample",
       y = expression(Log[2](Normalised~Precursor~Quantity))) +
  theme(axis.text.x = element_text(angle = 90))
```

```{r}
qcObj |>
  select("value", "CondRep","Drug","Pipeline") |>
  filter(!is.na(value)) |>
  ggplot() +
  aes(x = value,
      color = Drug,
      group = CondRep) +
  geom_density() +
facet_wrap(~Pipeline) +
  labs(title = "Normalised precursor quantity distributions per sample",
       x = expression(Log[2](Normalised~Precursor~Quantity))) +
  theme_bw()
```

```{r}
qcObj |>
  select("value", "Condition", "Replicate") |>
  filter(!is.na(value)) |>
  ggplot(aes(x = value,
           fill = Condition)) +
  geom_histogram() +
  scale_fill_observable() +
  facet_grid(Condition~Replicate) +
  labs(title = "Normalised precursor quantity distributions per sample",
       x = expression(Log[2](Normalised~Precursor~Quantity))) +
  theme_bw()
```

## Missed cleavages

QC on missed cleavage information.

```{r}
MC_alpha_mapping <- c("0" = 1,
                    "1" = 0.6,
                    "2" = 0.25)

qcObj |>
  select("value", "Replicate", "Condition", "Stripped.Sequence") |>
  filter(!is.na(value)) |>
    mutate(missed_cleavages = Stripped.Sequence |>
             str_count("[RK](?!(P|$))")) |> # calculate how many times an R or K appear in the Stripped.Sequence
  ggplot(aes(x = Replicate,
             fill = Condition)) +
  geom_bar(aes(alpha = missed_cleavages |> factor(levels = 4:0)),
           colour = "black") +
  scale_alpha_manual(values = MC_alpha_mapping, guide = guide_legend(override.aes = list(fill = pal_observable()(1)))) +
  scale_fill_observable() +
  facet_wrap(~Condition,
             scales = "free_x") +
  labs(title = "Missed cleavages per sample",
       x = "Replicate",
       alpha = "Missed cleavages") +
  theme_bw()
```

## Peptide types

### Trypticity

QC on peptide trypticity.

```{r}
qcObj |>
  select("value", "pep_type", "Replicate", "Condition") |>
  filter(!is.na(value)) |>
  mutate(pep_type = str_replace(pep_type, "-T", "-t")) |>
ggplot(aes(x = Replicate)) +
  geom_bar(aes(fill = pep_type),
           colour = "black"
           ) +
  scale_fill_nejm() +
  facet_wrap(~Condition,
             scales = "free_x") +
  labs(title = "Peptide types per sample",
       x = "Replicate",
       fill = "Peptide type") +
  theme_bw()
```

## Charge states

QC on charge states.

```{r}
qcObj |>
  select("value", "Precursor.Charge", "Replicate", "Condition") |>
  filter(!is.na(value)) |>
  filter(Precursor.Charge<=4) |>
ggplot(aes(x = Replicate)) +
  geom_bar(aes(fill = factor(Precursor.Charge, levels = 4:1)),
           colour = "black") +
  scale_fill_nejm() +
  facet_wrap(~Condition,
             scales = "free_x") +
  labs(title = "Peptide types per sample",
       x = "Replicate",
       fill = "Charge state") +
  theme_bw()
```

## MDS

Multi-dimensional scaling and PCA are equivalent if Euclidian distances are used.
The dist function, however, can handle missing values so the MDS implementation in R can handle missing values.

We make an MDS plot for each assay separately.

1. We convert the assay to a single cell experiment to use the runMDS function in the scater package
2. We make MDS plot

```{r}
se_lip <- runMDS(getWithColData(qf, "precursors_lip_norm") |> #.1
               as( "SingleCellExperiment"),
             exprs_values = 1)
plotMDS(se_lip, colour_by = "Condition") +
  geom_text_repel(aes(label = se_lip$CondRep),
                  size = 2,
                  show.legend = F) +
    labs(subtitle = "LiP assay (precursor level)") #.2
```
## Correlation matrix

```{r}
mat_lip <- assay(se_lip)
colnames(mat_lip) <- colData(se_lip)$CondRep
mat_lip |>
  assay() |>
  na.exclude() |>
  cor(method = "spearman") |>
  ggcorrplot::ggcorrplot() +
  scale_fill_viridis_c() +
  labs(title = "Correlation matrix",
       fill = "Correlation") +
  theme(axis.text.x = element_text(angle = 90))
```
Remove objects for plotting

```{r}
rm(list=c("se_lip", "mat_lip", "qcObj"))
gc()
gc()
```

# Statistics

We do a differential abundance analysis for the LiP data at the precursor level. 

## LiP Differential Analysis

The preprocessed data can now be modelled to infer the effect of the Drug.
The formula for the model thus becomes `formula = ~ Drug`
 
```{r, warning=FALSE}
qf <- msqrob(object = qf, 
             i = "precursors_lip_norm", 
             formula = ~ Drug)
```

We can now convert the research question "does the treatment affect the LiP precursor intensities?" into a statistical hypothesis. 

To aid defining the contrast, we will visualise the experimental design using the `ExploreModelMatrix` package. 


```{r}
colData_lip <- qf |> 
  getWithColData(i="precursors_lip_norm") |> 
  colData()

vd <- ExploreModelMatrix::VisualizeDesign(
    sampleData =  colData_lip,
    designFormula = ~ Drug,
    textSizeFitted = 4
)
vd$plotlist
```

We notice that the log2 FC between Rapa and DMSO is parameterised using the DrugRapa model parameter. 


1. We first define the contrast matrix, e.g. the linear combination of the model parameters that corresponds to the null hypothesis of interest, which is here defined as: 
`DrugRapa = 0`
2. We perform the hypothesis test for each precursor. 

```{r}
L <- makeContrast("DrugRapa=0", parameterNames = colnames(vd$designmatrix)) #1. 

qf <- hypothesisTest(object = qf, 
                     i = "precursors_lip_norm", 
                     contrast = L, 
                     overwrite = TRUE) #2. 
```

# Visualize results 

Extract results for lip assay

```{r}
contrast <- colnames(L)
results_lip <- rowData(qf[["precursors_lip_norm"]])[c("Precursor.Id", "Protein.Group", "Genes", "Proteotypic", "Stripped.Sequence", contrast)] |>
  data.frame() |>
  rename_with(~str_remove(.x,paste0(contrast,".")))
```


Set significance level at 5%

```{r}
alpha <- 0.05
```

## Differential abundance analysis on LiP

Significant hits based on differential abundance on LiP data.

```{r}
results_lip |>
  filter(adjPval <alpha) |>
  relocate(c("Precursor.Id","logFC","se","t","pval","adjPval")) |>
  arrange(pval) |> 
  knitr::kable()
``` 

```{r}
POI_DA <- results_lip |>
  filter(adjPval<alpha) |> 
  arrange(pval) |>
  pull("Protein.Group") |>
  unique()
POI_DA
```

## Volcano Plots 

We first make a helper function for a basic volcano plot. 

1. We calculate the adjusted significance level for the original p-values that corresponds to the nominal FDR significance level.
2. We make the volcano plot by plotting statistical significance, i.e. -log10(pvalues), in function of the effect size or biological relevance, i.e. log2 FC. We color the dots according to their significance, black if the adjusted p-value is higher than the FDR significance level and red if the adjusted p-value is lower than the FDR significance level. 
3. We add a line with the adjusted significance level that corresponds to the original p-values
4. We add the dots to the plot.
5. We set the layout of the plot (theme, labels, etc.)

```{r}
plot_volcano <- function(results_table, effectsize = "logFC", adjPval = "adjPval", pval = "pval", significance_level = 0.05, significance_colors= c("FALSE" = "black", "TRUE" = "red"), opacity = .5)
{

adjAlpha <- significance_level*mean(results_table[[adjPval]] < significance_level, na.rm = TRUE) #1.

volcano_out <- 
  results_table |> 
  filter(!is.na(.data[[adjPval]])) |> 
  ggplot( 
          aes(x = .data[[effectsize]],
              y = -log10(.data[[pval]]), col = .data[[adjPval]] < significance_level)) + #2.
  
  geom_hline(yintercept = -log10(adjAlpha)) + #3.
  
  geom_point(size = 1, alpha = opacity) + #4. 
  
  theme_bw() + #5. 
  ylab("-log10(p-value)") +
  scale_color_manual(values = significance_colors, breaks = c("TRUE", "FALSE")) +
  labs(color=paste0("FDR < ",significance_level))
return(volcano_out)
}
```

### Differential Analysis on LiP

```{r}
volcano_DA <- plot_volcano(results_lip)
volcano_DA
```

Now, we make volcano plot fancy. 

We first add the peptide type to the results 

1. Include mapping
2. Classify peptide 
3. Remove protein sequence and length

```{r}
results_lip <- results_lip |>     
  left_join(mapping, by = join_by(Protein.Group == Accession)) |> 
  mutate(pep_type = classify_trypticity(Stripped.Sequence, Protein.Sequence)) |>  
  select(-Protein.Length, -Protein.Sequence)
```

Set colors for proteins of interest to be plotted. 

```{r}
# POI_Plot <- POI_DA
# if (length(POI_Plot) > 10) POI_Plot <- POI_Plot[1:10]
# colors <- c(pal_npg()(length(POI_Plot)), "darkorange", "grey")
# names(colors) <- c(POI_Plot, "relevant", "non-relevant")

POI <- "P62942"
results_lip <- results_lip %>%
  mutate(Protein_type = case_when(
    Protein.Group == POI ~ "FKBP1A",
    adjPval < alpha & abs(logFC) > 1 ~ "Other",
    TRUE ~ "Not significant"
  ))
```

Make fancy volcanoplot

1. Remove geom_point layer
2. Add layer with all points
3. Add relevant log2 FC cutoff 
4. Add layer with relevant points (FDR < alpha and more extreme than log2 cutoff)
5. Add layer with POIs
6. Set colors shape and color
7. Change labels of shape and color legends

```{r}
fancy_volcano_DA <- delete_layers(volcano_DA,"GeomPoint") +
  theme(
    text = element_text(family = "Arial"),            # basislettertype
    axis.title = element_text(size = 14, face = "bold"),
    axis.text = element_text(size = 12),
    legend.title = element_text(size = 13, face = "bold"),
    legend.text = element_text(size = 12)
  ) +
  geom_point(shape=16, 
             color = "grey", 
             alpha=.2,
             data = results_lip |>
               filter(Protein_type == "Not significant")) +
  
  geom_vline(xintercept = c(-1,1), linetype = "dashed") +
  
  geom_point(aes(x=logFC, y = -log10(pval), color = Protein_type), 
             shape = 16,
             data = results_lip |> 
               filter(Protein_type %in% c("FKBP1A", "Other")), 
             alpha = .7,
             size = 2) +
  
  guides(shape = guide_legend(override.aes = list(alpha=1)),
         color = guide_legend(title = "Significant Proteins")) +
  scale_color_manual(values = c("FKBP1A" = "darkorange", "Other" = "black")) +
  #scale_shape_manual(values = c("Tryptic" = 16)) +
  #                    breaks = c("Ambiguous", "Tryptic"), # <-- Semi‑Tryptic verdwijnt uit de legende
  #                   name = "Peptide Type") +
  labs(x = expression(Log[2]*"(FC)"),
       y = expression(-Log[10]*"(p-value)"))
fancy_volcano_DA
```

Add gene-names for relevant genes and POIs.

```{r}
fancy_volcano_DA_genenames <- fancy_volcano_DA +
  theme(
    text = element_text(family = "Arial")) +
  ggrepel::geom_text_repel(
    aes(x = logFC, y = -log10(pval), label = Genes),
    color = "darkorange",
    size = 3,
    segment.size = 0.25,
    show.legend = F,
    data = results_lip |> 
      filter(Protein.Group == POI & adjPval < alpha & abs(logFC) > 1 & !is.na(pval))
    )
  # ggrepel::geom_text_repel(
  #   data = results_lip %>% 
  #     filter(Protein.Group != POI & adjPval < alpha & abs(logFC) > 1 & !is.na(pval)),
  #   aes(x = logFC, y = -log10(pval), label = Genes),
  #   color = "black",
  #   size = 3,
  #   segment.size = 0.25,
  #   show.legend = FALSE
  # )
x_max <- ceiling(max(abs(results_lip$logFC), na.rm = TRUE))
fancy_volcano_DA_genenames <- fancy_volcano_DA_genenames +
  scale_x_continuous(
    limits = c(-x_max, x_max),               # 0 exact in het midden
    breaks = scales::breaks_extended(5),     # nette as-stappen
    expand = expansion(mult = c(0.02, 0.02)) # klein randje links/rechts
  )
fancy_volcano_DA_genenames

volcano_imput <- fancy_volcano_DA_genenames
saveRDS(volcano_imput, "volcano_imput.rds")

ggsave(
  filename = "../output/volcano_PELSA_fancy_imput.png",
  plot = fancy_volcano_DA_genenames,
  width = 8,
  height = 4,
  dpi = 600
)
```

## Stuctural Barcode plots 

We will process the results table by 

1. Unnesting `Protein.Groups` containing more than one protein and we will introducing new variables accession and gene that contains just a single protein and gene. 
2. Map the accessions in the results table to the accessions in the mapping table
3. Retrieve all start and end positions for each peptide in the protein (and thus multiple start positions for peptides that are repeated within a protein)
4. str_locate_all returned a list for each peptide with all start and end positions. We use map from the purrr packages to obtain a list with the start positions only. 
5. We unnest start positions for repeated peptide sequences.
6. We determine the peptide type.
7. We convert the table to a data.frame

```{r}
results_lip_barcode <- results_lip |>
  mutate(accession = strsplit(Protein.Group, ";"),
         gene = strsplit(Genes, ";")) |> #1.
  unnest(c(accession, gene)) |> #1.
  left_join(mapping, by = join_by(accession == Accession)) |> #2.
  mutate(tmp = str_locate_all(Protein.Sequence, Stripped.Sequence), #3.
         start = purrr::map(tmp, ~ .x[, "start"]),
         end = purrr::map(tmp, ~ .x[, "end"])) |> #4.
  unnest(c(start, end)) |> #5.
  mutate(
         pep_type = classify_trypticity(peptide = Stripped.Sequence, protein = Protein.Sequence, start_pos = start)) |> #6.
  as.data.frame() #7. 
```

We calculate the observed intensities in each group and add them to the results matrix.

```{r}
se <- getWithColData(qf, "precursors_lip_norm")
groupsMatrix <- model.matrix(~-1 + Drug, data = colData(se))
nobs_group <- ((!is.na(assay(se))) %*% groupsMatrix) |> 
  as.data.frame() |>
  tibble::rownames_to_column("Precursor.Id")
results_lip_barcode  <- results_lip_barcode  |> 
  left_join(nobs_group, by = "Precursor.Id") 
```

```{r}
results_lip_barcode <- results_lip_barcode |> mutate( 
  missing = do.call(pmin,pick(contains("Drug")))==0,
  significant = ifelse(missing, 
                       "missing", 
                       ifelse(
                         is.na(adjPval),
                         NA, 
                         ifelse(adjPval < 0.05, 
                                "significant",
                                "non-significant")
                         )
                       ) |> factor(levels=c("non-significant","missing", "significant")),
  directionality = ifelse(is.na(significant), 
                          NA, 
                          ifelse(significant == "significant",
                                  ifelse(logFC < 0, 
                                         "up_DMSO", 
                                         "up_Rapa"), 
                                  ifelse(significant == "missing",
                                         ifelse(DrugDMSO==0,
                                                "missing_DMSO",
                                                "missing_Rapa"),
                                         "non-significant")
                                  )
                          )
)
```

### Barcode plots

We first make a function for the barcode plots
1. Checks if all levels of the color_variable are matched by the names of the colors
2. Checks if the end of all peptide sequences is smaller than the protein length 
3. It adds an empty plot
4. Uses manual fill values defined by colors
5. Defines the theme of the plot
6. Loops over all colors names in variable `color` in reverse order and adds the segments with the corresponding color in the variable `color_variable`
7. Uses the length of the protein to specify the limits of the x-axis if `protein_length` is provided.
8. Generates a title if the `protein_length`, `protein_name` or `gene_name` is specified. 


```{r}
barcode_plot <- function(poi_data, color_variable, colors, protein_length = NULL, protein_name = NULL, gene_name=NULL)
{
  if(sum(!(unique(poi_data[[color_variable]]) %in% names(colors))) > 0) stop(paste0("color_variable:",  color_variable," contains levels in poi_data object for which no colors are provided in argument colors")) #1.
  
  if(!is.null(protein_length)) if(protein_length < max(poi_data$end)) stop(paste0("Some features map beyond protein length: ",max(poi_data$end)," > ", protein_length)) #2.
  
  barcode_plot <- poi_data |> 
    ggplot(aes(x=start, y = 1))  + #3.
    scale_fill_manual(values = colors, breaks = names(colors)) + #4.
    theme_bw() + theme(panel.grid = element_blank(), #5.
          axis.line.y = element_blank(),
          axis.text.y = element_blank(),
          axis.ticks.y = element_blank(),
          panel.background = element_rect(fill = "white"),
          strip.background = element_rect(fill = "white"),
          strip.text = element_text(face = "bold"))
  
  for (type in names(colors)[length(colors):1]) #6. 
    if (type %in% poi_data[[color_variable]])
      barcode_plot <- barcode_plot + geom_rect(aes(xmin = start-1,
                  xmax = end,
                  ymin = .5,
                  ymax = 1.5,
                  fill = .data[[color_variable]]
                  ),
              linewidth = 0.2, 
              data=poi_data |> 
                filter(poi_data[[color_variable]]==type)
              )
  
    if(!is.null(protein_length)) barcode_plot <- barcode_plot + coord_cartesian(
  xlim = c(0, protein_length), expand = FALSE) #7. 
  
    if(!is.null(protein_length) | !is.null(protein_name) | !is.null(gene_name)) #8.
    {
      subtitle <- paste0(protein_name, 
                         ifelse(is.null(protein_name) | is.null(gene_name),"","/"), 
                         gene_name, 
                         ifelse(!is.null(protein_name) | !is.null(gene_name) & !is.null(protein_length),": ", ""),
                         round(calculate_coverage(
                           poi_data$start,
                           poi_data$end,
                           protein_length)*100,
                           1), 
                         "% coverage"
                         )
    barcode_plot <- barcode_plot +   labs(subtitle = subtitle)
    }
  return(barcode_plot)
}
```

#### Differential Abundance

We will loop over all proteins of interest and make an barcode plot.

1. We set colors for the plot. 
2. We make a list to store the plot objects for the different POI
3. We loop over the POI 
4. We store a barcode plot in for each poi
5. We select all features belonging to the poi
6. We color the features according to the `significant` variable in the filtered results_lip_barcode object
7. We use our custom defined colors
8. We define the protein_name
9. We retrieve the protein length
10. We retrieve the gene_name
11. We include the plot in the vignette output

```{r}
colors <- c("significant" = "orange", "missing" = "blue", "non-significant" = "grey40") #1.
da_barcode <- list() #2.
for (poi in POI_DA){ #3.
  da_barcode[[poi]] <- 
    barcode_plot(results_lip_barcode |> #4
                   filter(accession==poi & !is.na(significant)), #5
                 color_variable = "significant", #6
                 colors = colors, #7.
                 protein_name = poi, #8.
                 protein_length = mapping |> #9.
                   filter(Accession == poi) |>
                   pull(Protein.Length) |> 
                   unlist(),
                 gene_name = results_lip_barcode |>  #10.
                   filter(accession==poi) |> 
                   pull(gene)|>
                   unique()) +
    scale_fill_manual(
    values = colors,
    breaks = names(colors),
    labels = c(
      "significant" = "Significant",
      "missing" = "Missing",
      "non-significant" = "Non-significant"
    )
  ) +
    theme(
      text = element_text(family = "Arial"),  # Arial
      axis.title.y = element_blank(),          # geen y-as label
      axis.text.y = element_blank(),           # geen y-as ticks
      axis.ticks.y = element_blank(),          # geen y-as ticks
      plot.subtitle = element_text(size = 10, face = "bold"),
      axis.title.x = element_text(size = 8, face = "bold"),
      axis.text.x = element_text(size = 6, color = "black"),
      legend.title = element_text(size = 10, face = "bold"),
      legend.text = element_text(size = 8)
  ) +
    labs(
    x = "Protein Sequence",     # X-as label
    fill = "Significance"        # Legend titel voor geom_rect(fill = ...)
  )
                       
  print(da_barcode[[poi]]) #11.  
}

ggsave(
  filename = "../output/barcode_FKBP1A_simple_imput.png",
  plot = da_barcode[[4]], # nr.4 = FKBP1A
  width = 6,
  height = 1.5,
  dpi = 600
)
```

With directionality

```{r}
colors <- c("up_DMSO"="lightslateblue", "up_Rapa" =   "orange2", "missing_DMSO" =   "yellow2", "missing_Rapa" = "lightskyblue", "non-significant" = "grey40")
da_barcode_directionality <- list() #2.
for (poi in POI_DA){ #3.
  da_barcode_directionality[[poi]] <- 
    barcode_plot(results_lip_barcode |> #4
                         filter(accession==poi & !is.na(significant)), #5
                 color_variable = "directionality", #6
                 colors = colors, #7.
                 protein_name = poi, #8.
                 protein_length = mapping |> 
                   filter(Accession == poi) |>
                   pull(Protein.Length) |> 
                   unlist(), #9.
                 gene_name = results_lip_barcode |> 
                   filter(accession==poi) |> 
                   pull(gene)|>
                   unique()) + #10.
    scale_fill_manual(
    values = colors,
    breaks = names(colors),
    labels = c(
      "up_DMSO" = "Up DMSO",
      "up_Rapa" = "Up Rapa",
      "missing_DMSO" = "Missing DMSO",
      "missing_Rapa" = "Missing Rapa",
      "non-significant" = "Non-significant"
    )
  ) +
  theme(
      text = element_text(family = "Arial"),  # Arial
      axis.title.y = element_blank(),          # geen y-as label
      axis.text.y = element_blank(),           # geen y-as ticks
      axis.ticks.y = element_blank(),          # geen y-as ticks
      #plot.subtitle = element_blank(),
      plot.subtitle = element_text(size = 10, face = "bold"),
      #axis.title.x = element_blank(),
      axis.title.x = element_text(size = 8, face = "bold"),
      axis.text.x = element_text(size = 8, color = "black"),
      legend.title = element_text(size = 10, face = "bold"),
      legend.text = element_text(size = 8)
  ) +
    labs(
    x = "Protein Sequence",     # X-as label
    fill = "Directionality"        # Legend titel voor geom_rect(fill = ...)
  )
  print(da_barcode_directionality[[poi]]) #11.  
}

fkbp_imput <- da_barcode_directionality[[4]]
saveRDS(fkbp_imput, "fkbp_imput.rds")

ggsave(
  filename = "../output/barcode_FKBP1A_dir_imput.png",
  plot = da_barcode_directionality[[4]], # nr.4 = FKBP1A
  width = 6,
  height = 1.5,
  dpi = 600
)
```


# Session Info

For reproducibility, we include a session info in our script so readers can see our particular setup of R.

```{r}
sessionInfo()
```
````
