## Supplementary material for "Experimental mismatch in benchmarking PELSA and LiP-MS": README.txt

### README — DIA-LiPA Reanalysis ###

This repository contains the full analysis structure used for the DIA-LiPA reanalysis of the publicly available PELSA rapamycin dataset. 
All files required to reproduce the analysis from processed Spectronaut output are provided.

# Repository Structure
/Imputed_data
    /input
        SampleAnnotation.txt
        SP_9606.fasta
        Spectronaut_report.tsv
    /output
        barcode_FKBP1A_dir.png
        fkbp.rds
        volcano_PELSA_fancy.png
        volcano.rds
    /script
        DIA-LiPA-unpaired.R

/NotImputed_data
    (identical structure as above)

# Folder Descriptions
Imputed_data and NotImputed_data

Each folder contains a complete analysis workflow. The only difference is whether the data were imputed prior to analysis.

input/
SampleAnnotation.csv
-> Metadata file describing raw file names, conditions and replicates used for statistical testing.

SP_9606.fasta
-> FASTA database used for peptide/protein annotation.

Spectronaut_report.tsv
-> Direct output from Spectronaut used as input for the R script.

script/
DIA-LiPA_unpaired.R
-> R script that processes the input files and generates all analysis outputs.

output/
barcode_FKBP1A_dir.png — structural barcode for FKBP1A with directionality
volcano_PELSA_fancy.png — relative differential abundance volcano plot rapamycin versus DMSO


# How to Reproduce the Analysis
1) Open R or RStudio.
2) Set the working directory to either Imputed_data/script or NotImputed_data/script.
3) Run the script.
4) All generated figures will appear in the corresponding output/ folder.

# Notes
Raw files are not included because this is a reanalysis based on already published datasets.
The Spectronaut output distributed here is sufficient to reproduce all results demonstrated in the manuscript.
