## Supplementary figures and images for "Experimental mismatch in benchmarking PELSA and LiP-MS"

### barcode_FKBP1A_dir_imput.png

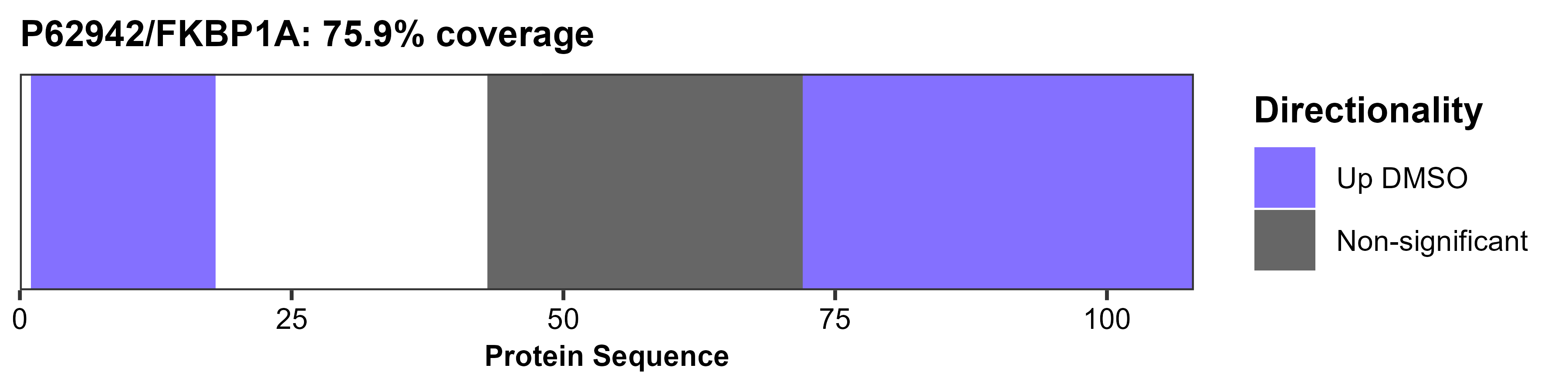

### barcode_FKBP1A_dir_noimput.png

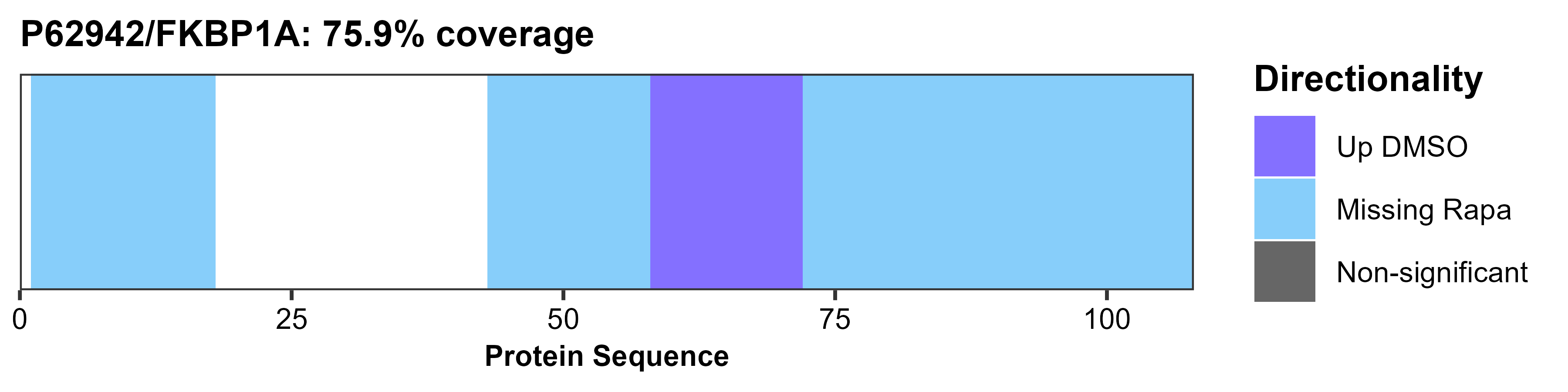

### volcano_PELSA_fancy_imput.png

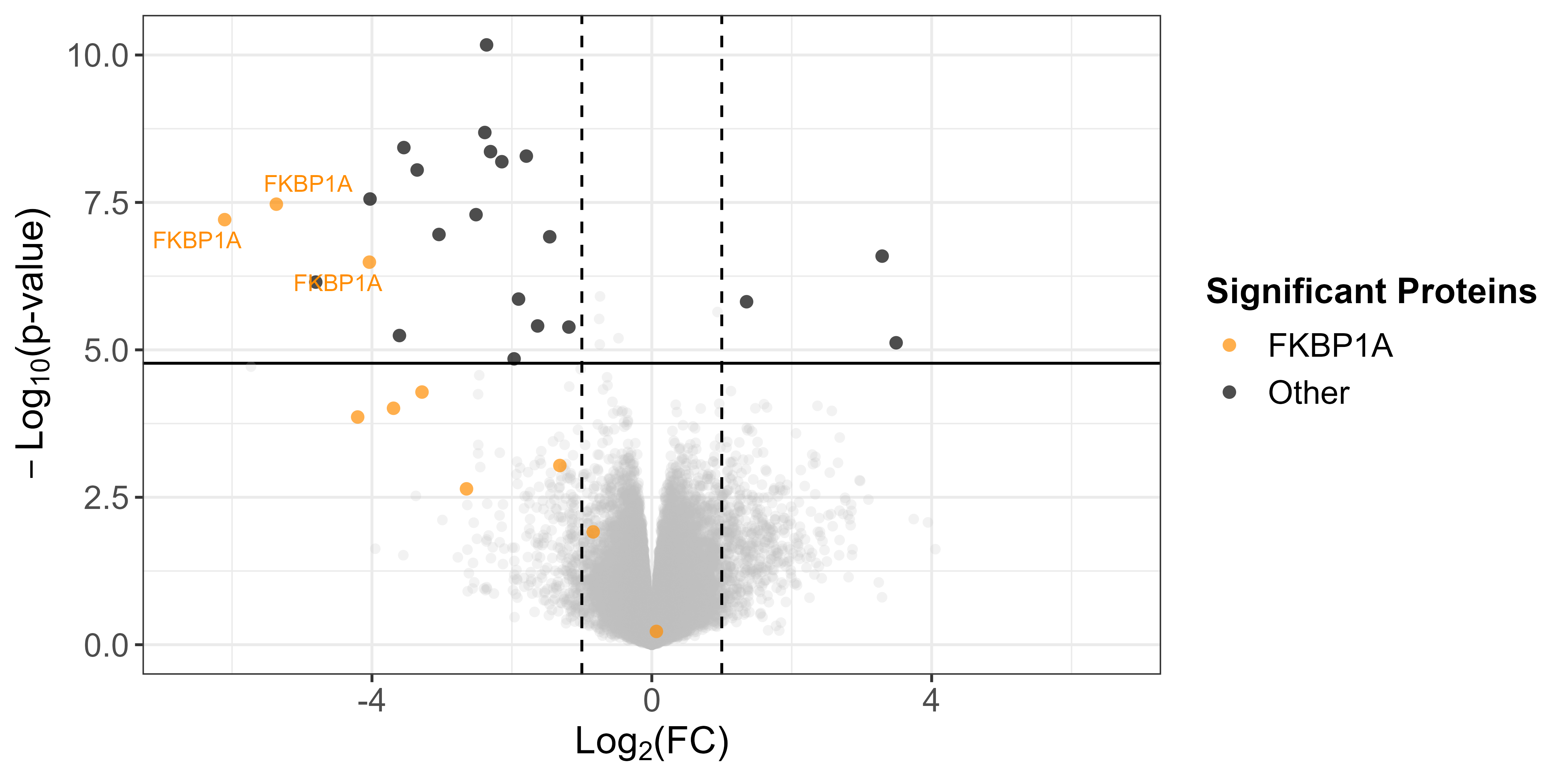

### volcano_PELSA_fancy_noimput.png

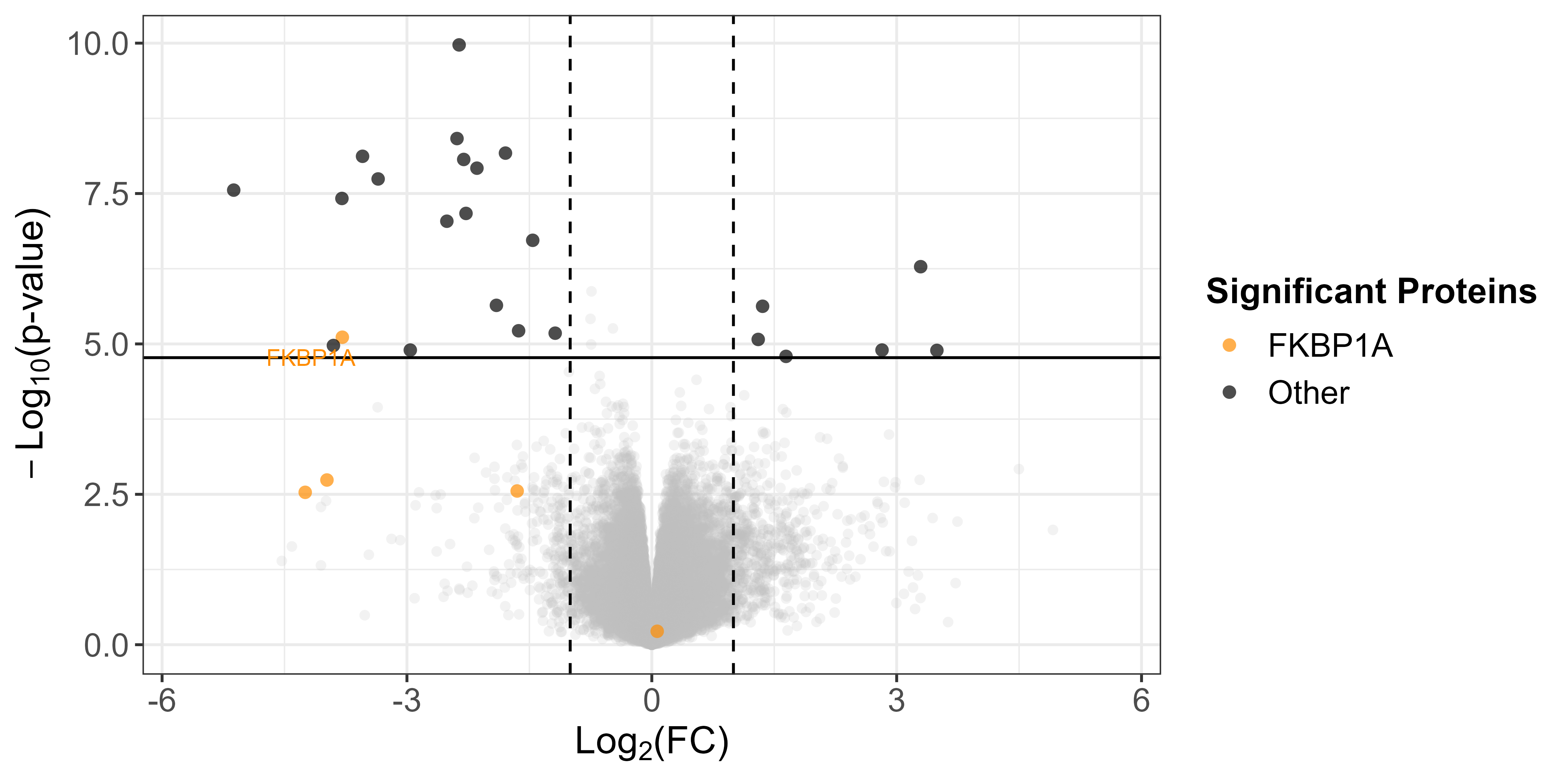
